# Thyroid cancer incidence and mortality in Latin America

**DOI:** 10.1101/487249

**Authors:** Fábia Cheyenne Gomes de Morais Fernandes, Dyego Leandro Bezerra de Souza, Maria Paula Curado, Isabelle Ribeiro Barbosa

## Abstract

This study analyzed trends in thyroid cancer incidence and mortality in countries of Latin America. Ecological study of time series, with incidence data extracted from the International Agency for Research on Cancer (IARC), in the 1990-2012 period and mortality data obtained from 16 countries of the World Health Organization (WHO), in the 1995-2013 period. The trend of incidence rate was analyzed by the Joinpoint regression. The average annual percentage change (AAPC) and the 95% confidence interval (CI 95%) were calculated for incidence and mortality. The average rate of thyroid cancer incidence was higher in Quito (Ecuador) between the ages of 40 to 59 years old, 42.2 new cases per 100,000 inhabitants, as well as mortality 4.8 deaths per 100,000 women inhabitants above 60 years old. There was an increase in thyroid cancer incidence trends in women, for all age groups, in Cali, Costa Rica and Quito and men in Costa Rica; there was stability above the age of 60 years old in Cali, Goiania, Quito and Valdivia in men, as well as women in Goiania and Valdivia. There was a trend of increasing mortality for females in three countries: Ecuador (AAPC= 3,28 CI 95% 1,36;5,24), Guatemala (AAPC= 6,14 CI 95% 2,81;9,58) and Mexico (AAPC= 0,67 CI 95% 0,16;1,18). Thyroid cancer in Latin America showed a high incidence, with increased incidence in women. Stability in mortality was observed for most countries of Latin America.

## INTRODUCTION

The thyroid malignancy neoplasm represents approximately 2% of all cancers in the world (1). In 2016, were estimated 238,000 new cases of thyroid cancer and 43.000 deaths for both sexes in the world, with a standardized incidence rate of 2.2/100,000 for male inhabitants and 4.4/100.000 for female inhabitants female (2). The incidence rate in Northern Americans exceeds most European countries (3, 4).

In South America, in 2012, the incidence rate for thyroid cancer was 8.4 cases/100.00 women and 1.9/100,000 men (1). It was classified between the fifth to the tenth most common type of cancer among women and the 25th most commonly diagnosed cancer in men in Central and South America (5). There is an increased tendency to the female incidence in Brazil, Colombia, Costa Rica, and Ecuador, as the males in Brazil and Costa Rica (6). High income countries show a substantial increase in the incidence (7), considered to be two times higher compared to low/medium income countries, for both sexes (8).

Thyroid cancer is considered rare, but in the last 30 a sharp increase in incidence was registered, attributed to higher diagnostic intensity, although environmental influences, genetic and dietary merit further investigation (9, 10). This cancer is characterized with predominance in females, white race and the median age of 45 years, but with a tendency to increase among young adults (9, 11, 12).

Latin America faces difficulties to fulfill an agenda of continuous commitments, with appropriate financial incentives and effective approaches to cancer control. Therefore it becomes relevant to describe the epidemiological profile of thyroid cancer in these countries. The aim of this study was analyze the incidence, mortality, rates, and trends for thyroid cancer in Latin America.

## METHODS

This is about an ecological time series study, based on secondary data available in the databases of the International Agency for Research on Cancer (IARC) and the World Health Organization (WHO) (13,14). Occurrence and mortality for malignant thyroid neoplasm were analyzed in Latin American countries.

Incident cases of thyroid malignancy, during the period of 22 years (1990-2012), were extracted from the *Cancer Incidence in Five Continents* - CI5 PLUS, which includes five Population Based Cancer Registries (PBCRs): four regional registries, Cali (Colombia), Quito (Ecuador), Goiania (Brazil) and Valdivia (Chile) and one national record, Costa Rica (13). The mortality data, for which the available information was analyzed of 16 countries of Latin America, accounted for 90% of the population in Latin America between 1995 to 2013 (14).

The number of cases was extracted and adjusted rate-specific by age and calculated for three age groups (25-39, 40-59, and 60-74) and for all ages. Specific rates, adjusted by age, were calculated using the standard world population, according to sex and countries with available data.

The incidence rates and standardized mortality rates were calculated by sex. Also were calculated the Ratio of incidence and mortality rates by sex. The average annual percentage change (AAPC) was estimated for incidence and mortality with a 95% confidence interval (CI 95%) in the period. The exception to these analyses were Suriname and Uruguay, due to the lack of cases in the historical series. The statistical analyses were performed using the software *Joinpoint Regression*, version 4.5.0.0 (15,16).

## RESULTS

Between 1990 and 2012, the higher incidence rates of thyroid cancer were observed in Quito (Ecuador) and Costa Rica, in females, aged 40 to 59 years, with rates of 41.2/100,000 to 28.0/100,000, respectively. For males, the highest rates were observed in Quito (Ecuador) and Cali (Colombia), over 60 years, 11.7/100,000 to 7.3/100,000, respectively. (Table 1, Figure 1).

**Table 1.**
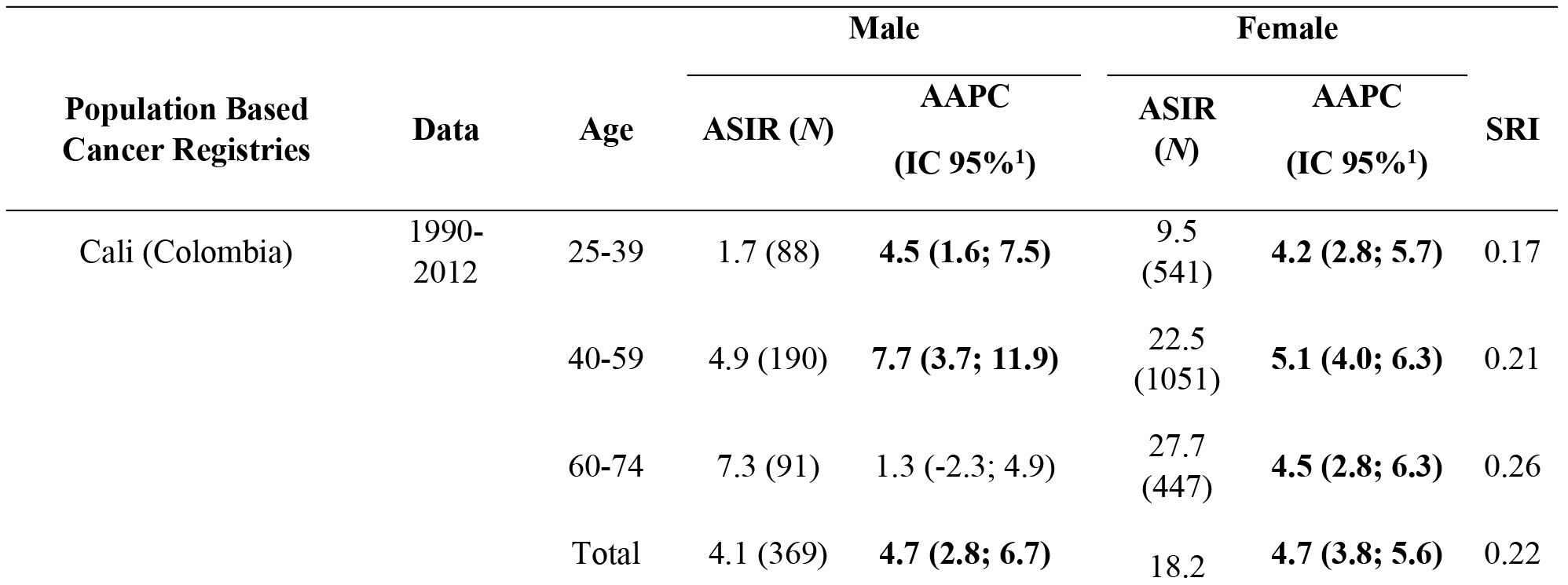
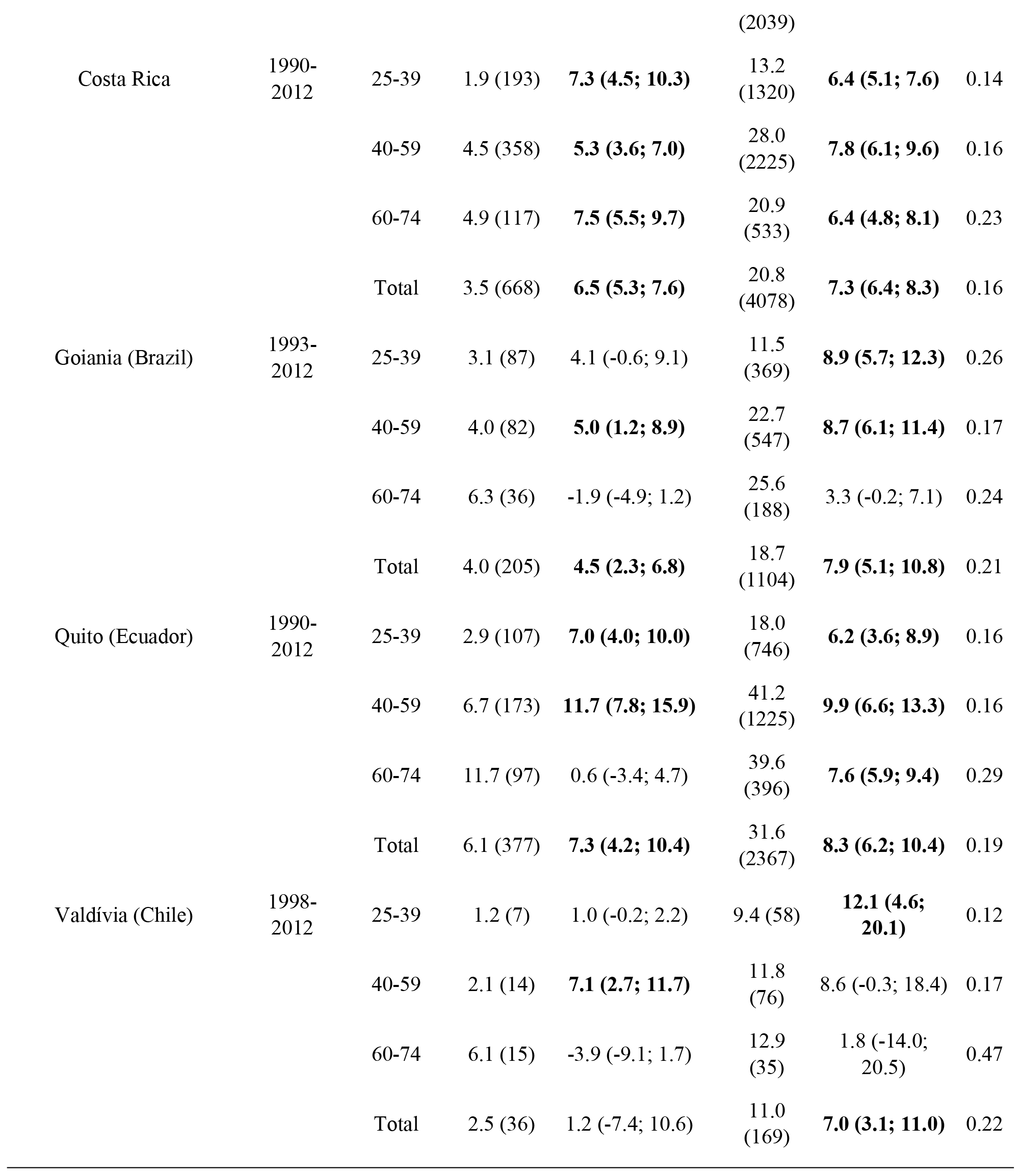
Age standardized incidence rate (ASIR), number of cases (*N*), average annual percent change (AAPC), and incidence rate ratio (SIR) for thyroid cancer, according to age and sex, in Cali (Colombia), Costa Rica, Goiania (Brazil), Quito (Ecuador) and Valdivia (Chile), for the period 1990–2012.

**Figure 1.**
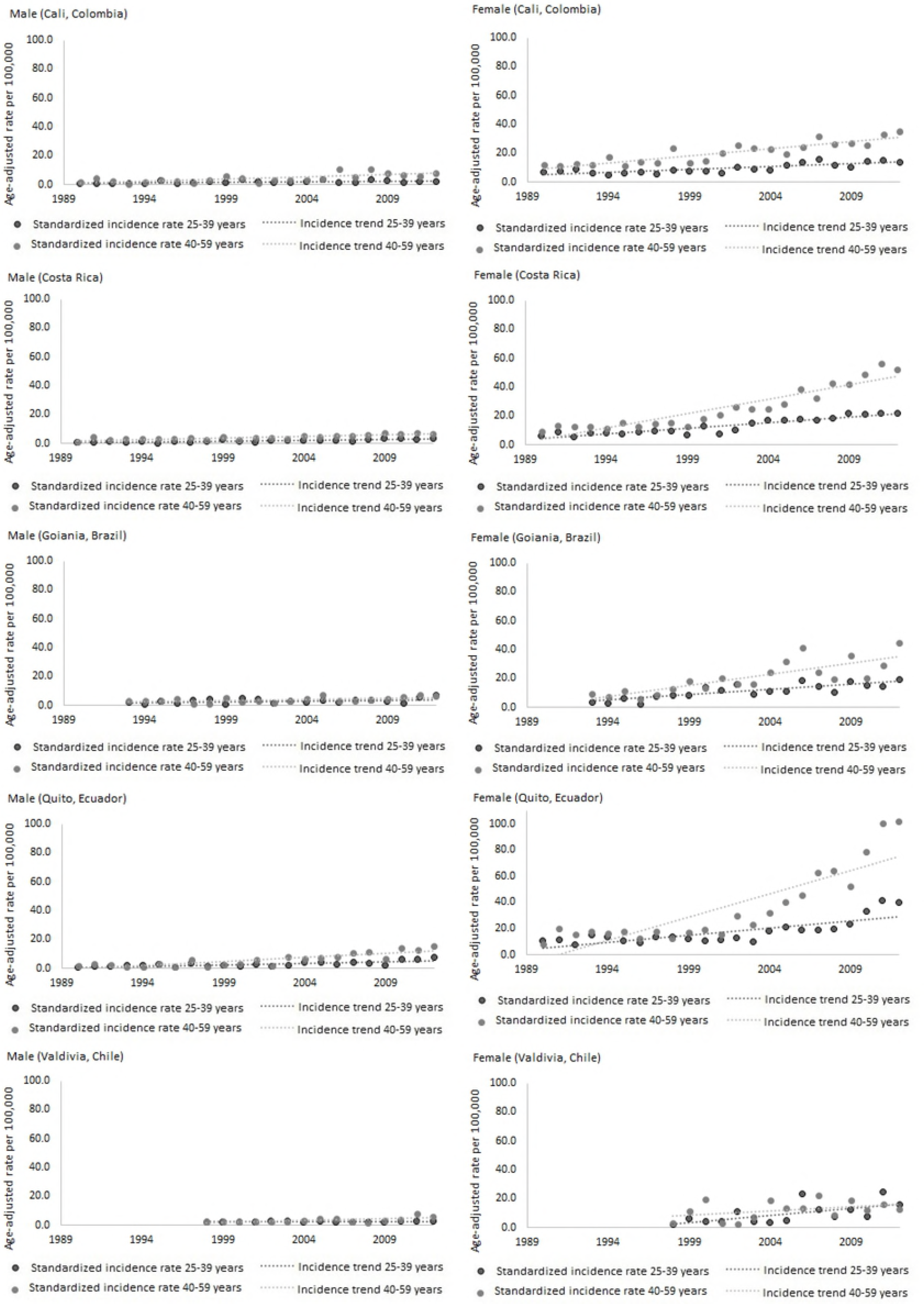
Thyroid cancer age-adjusted incidence rates (95% confidence interval) for sex, age range 25-39 and 40-59 years, for Cali (Colombia), Costa Rica, Goiania (Brazil), Quito (Ecuador) and Valdivia (Chile), for the period 1990–2012. 95% CI: 95% confidence interval. The grayline represents trends over the period.

Increasing incidence trends for thyroid cancer were evident in both sexes and all age groups, with the exception of the males, of Group 60 to 74 years in Cali (Colombia) 1.3% [95% CI:−2.3; 4.9]; Goiania (Brazil), between 25-39 years [4.1%; CI 95%: −0.6; 9.1] and between 60-74 years old [−1,9%; CI 95%: −4,9; 1,2]; Quito (Ecuador), between 60-74 years [0.6%; CI 95%:−3.4; 4.7]; Valdivia (Chile), between 25-39 years [1.0%; CI 95%:−0.2; 2.2) and between 60-74 years [−3.9%; CI 95%:−9.1; 1.7]. For females, there was stability in the age group between 60-74 years in Goiania (Brazil), [3.3%; CI 95%:−0.2; −7.1) and Valdivia (Chile), between 40-59 years [8.6%; CI 95%:−0.3; 18.4] and between 60-74 years [1.8%; CI 95%:−14.0; 20.5]. The incidence trends by age group followed similar patterns in Cali (Colombia), Costa Rica, Goiania (Brazil), Quito (Ecuador) and Valdivia (Chile), with the highest increase trends for women (Table 1, Figures 1, 2 and 3.).

**Figure 2.**
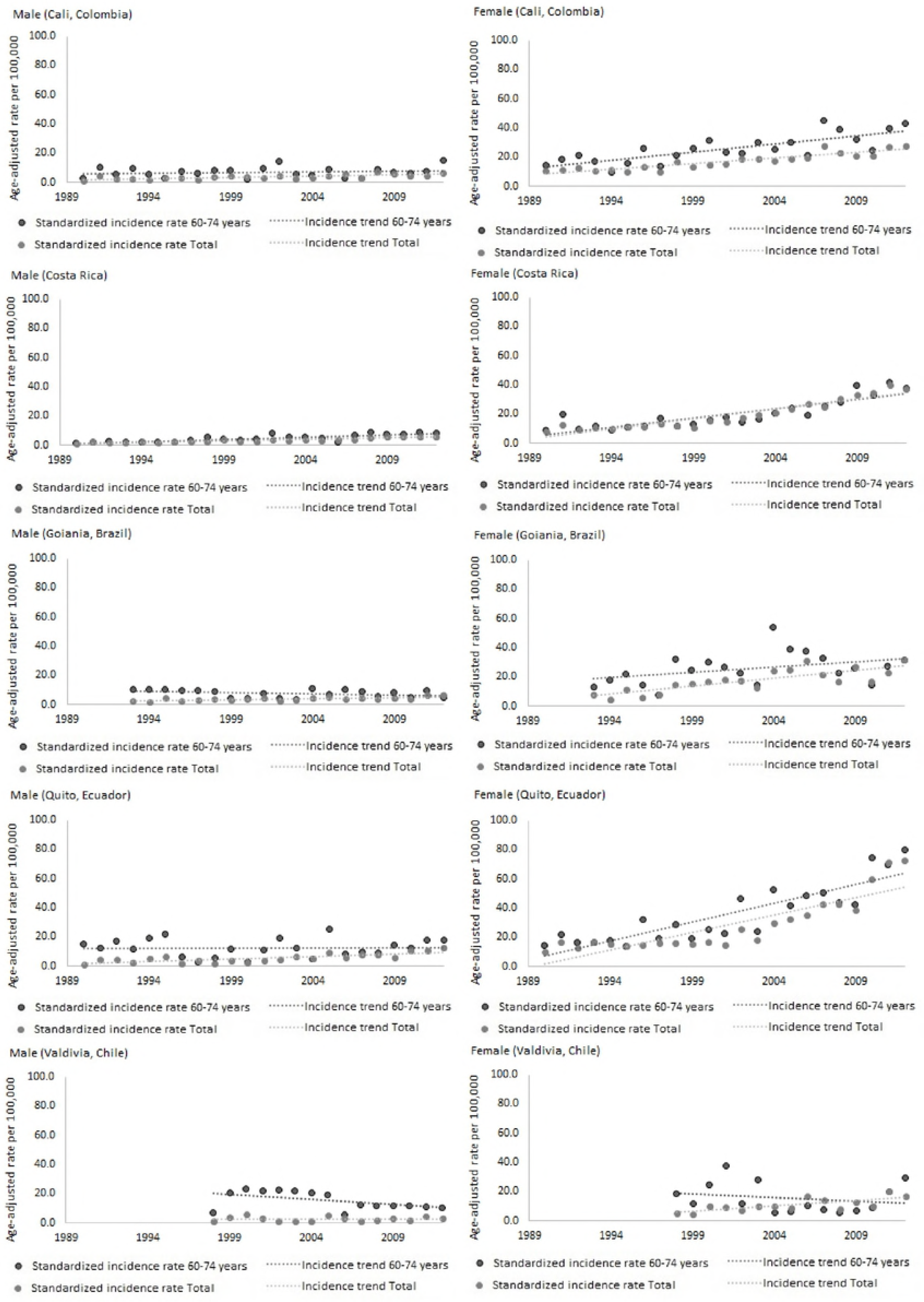
Thyroid cancer age-adjusted incidence rates (95% confidence interval), by sex, age 60-74 years and total, for Cali (Colombia), Costa Rica, Goiania (Brazil), Quito (Ecuador) and Valdivia (Chile), for the period 1990–2012. 95% CI: 95% confidence interval. The grayline represents trends over the period.

**Figure 3.**
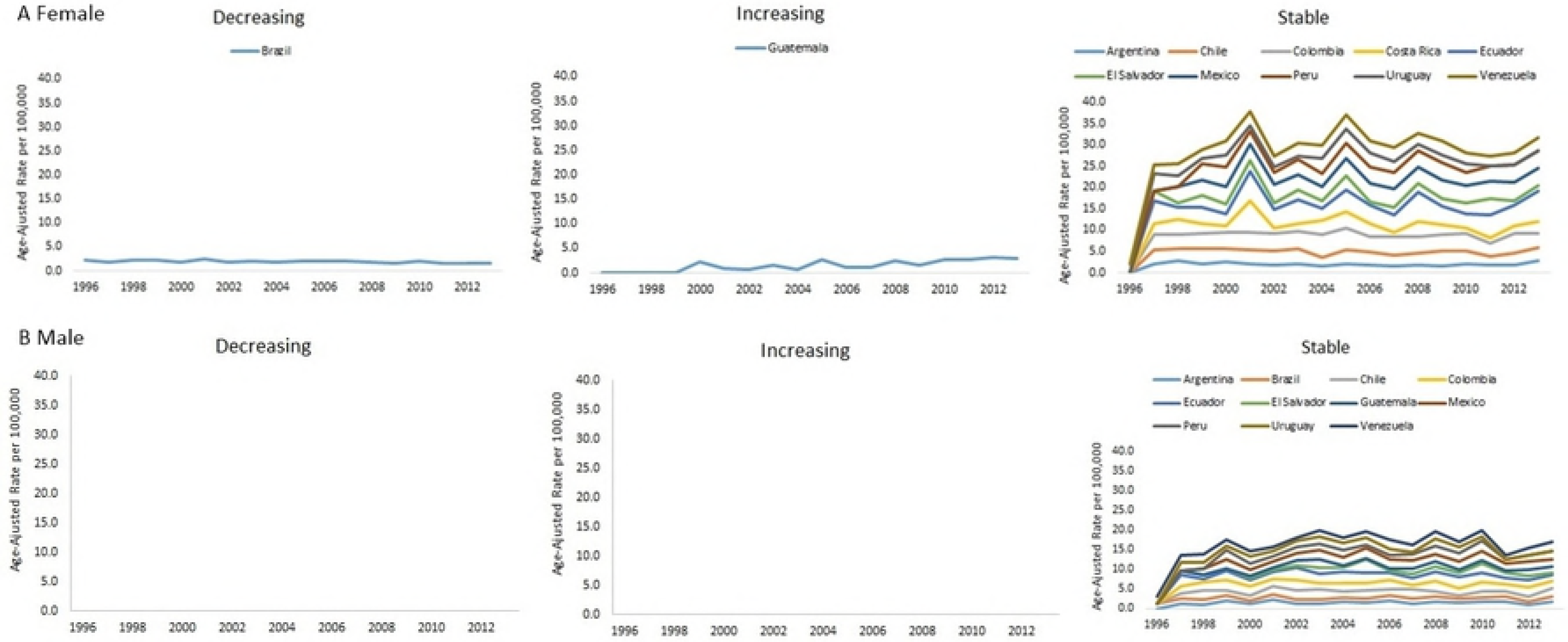
Temporal trends in thyroid cancer mortality, according to sex, and age 60-74 years, in 16 Latin American countries, for the period 1995–2013. (A) Female; (B) Male.

Between 1995 and 2013, the highest rates of thyroid cancer mortality were observed in Ecuador, in the age group 60-74 years (2.2/100,000 in men and 4.8/100,000 women). According to the ratio of the rates (ASMR), mortality was higher in women in most countries but presented Ratio equal to 1 in Ecuador between 25-39 years and Uruguay in groups of 25-39 years and 40-59 years (Table 2).

**Table 2.**
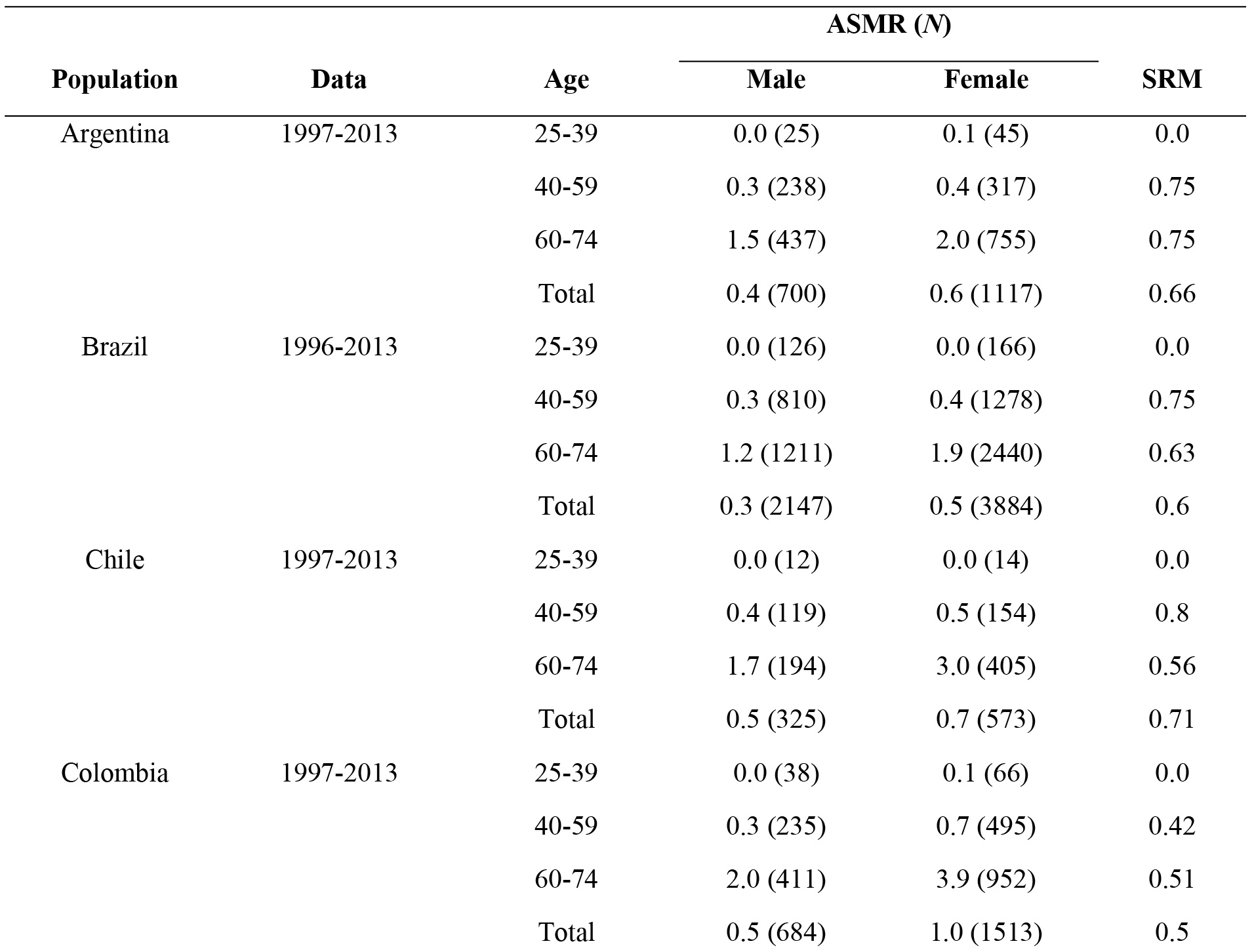
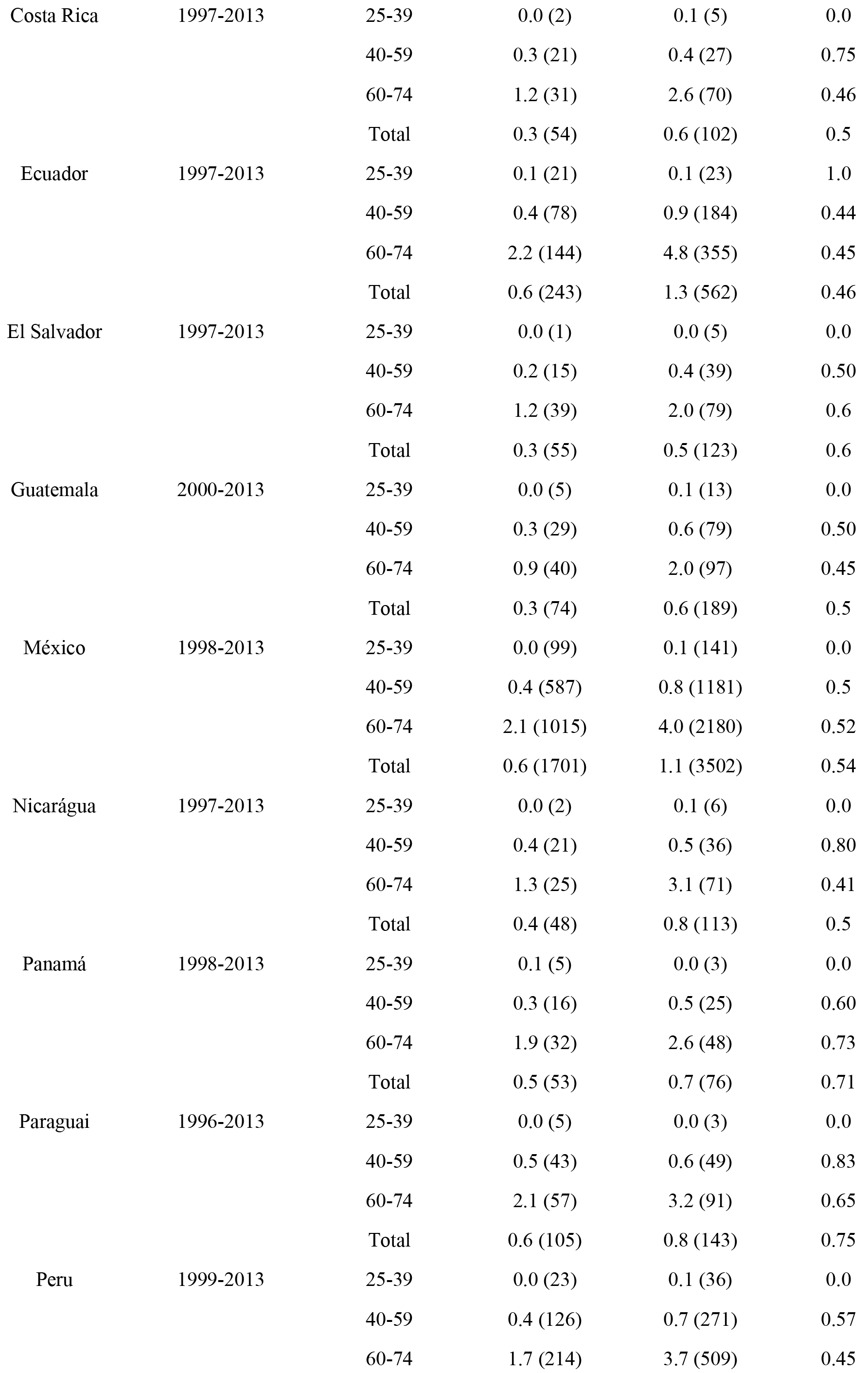
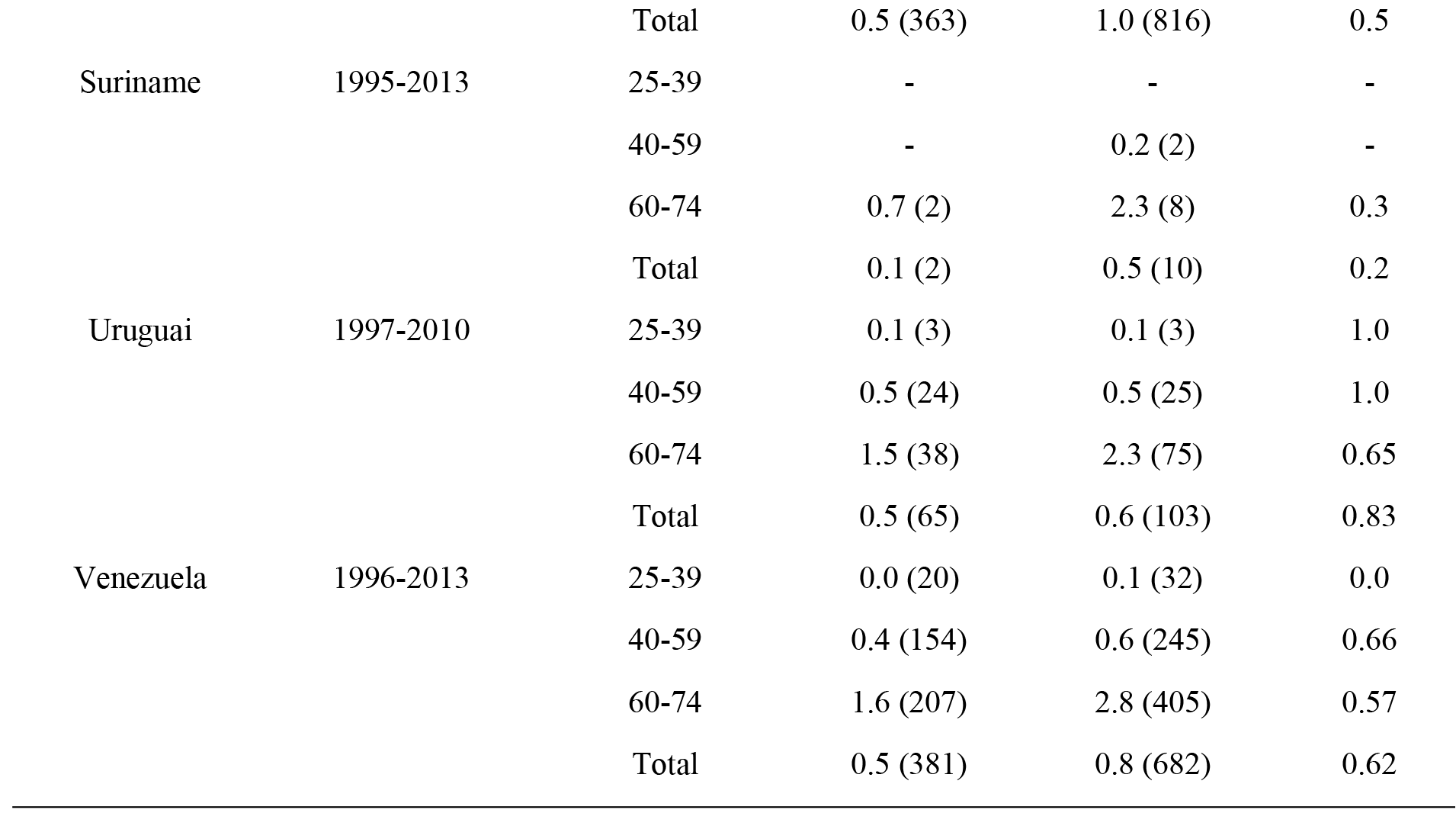
Age standardized mortality rate (ASMR) per 100,000, number (*N*) of deaths and mortality rate ratio (SMR) for thyroid cancer, by sex and age-group, for 17 Latin American populations, in the period 1995-2013.

There was no homogeneous pattern in trends of mortality in the 16 countries studied (Figure 4). In the age group 60-74 years, for women, it was observed an increase in Guatemala and reduction in Brazil. For men, the eleven countries that had historical series showed a trend of stability (Argentina, Brazil, Chile, Colombia, Ecuador, El Salvador, Guatemala, Mexico, Peru, Uruguay,and Venezuela). There was reduction in male mortality rate in Brazil, aged 40-59 years (−1.23%; CI 95%:-2.42; -0.02), while in female, the highest increase was observed in Guatemala (8.77%; CI 95%: 3.06-14.81), for all ages (Table 3).

**Table 3.** Thyroid cancer mortality trends, by sex and age-group for 17 Latin American populations, in theperiod 1995–2013.

| Population | Data | Age | AAPC (95% CI) |  |
| --- | --- | --- | --- | --- |
|  |  |  | Male | Female |
| Argentina | 1997-2013 | 25-39 | - | - |
|  |  | 40-59 | -1.11 (-4.18; 2.07) | 0.52 (-1.81; 2.90) |
|  |  | 60-74 | 0.70 (-2.03; 3.51) | -1.06 (-2.73; 0.64) |
|  |  | Total | 0.26 (-1.70; 2.26) | -0.62 (-1.88; 0.66) |
| Brazil | 1996-2013 | 25-39 | 0.29 (-3.79; 4.55) | 0.86 (-3.51; 5.44) |
|  |  | 40-59 | -1.23 (-2.42; -0.02) | -0.76 (-1.69; 0.18) |
|  |  | 60-74 | -0.37 (-1.63; 0.90) | -1.46 (-2.33; -0.59) |
|  |  | Total | -0.62 (-1.48; 0.24) | -1.15 (-1.81; -0.49) |
| Chile | 1997-2013 | 25-39 | - | - |
|  |  | 40-59 | -3.14 (-7.18; 1.09) | -1.42 (-4.98; 2.27) |
|  |  | 60-74 | -1.00 (-3.99; 2.09) | -0.91 (-2.50; 0.71) |
|  |  | Total | -1.95 (-4.01; 0.16) | -0.97 (-2.43; 0.50) |
| Colombia | 1997-2013 | 25-39 | - | - |
|  |  | 40-59 | -0.26 (-3.00; 2.56) | 0.52 (-1.60; 2.67) |
|  |  | 60-74 | -0.94 (-3.02; 1.19) | 0.11 (-1.38; 1.61) |
|  |  | Total | -0.84 (-2.27; 0.62) | 0.35 (-0.92; 1.63) |
| Costa Rica | 1997-2013 | 25-39 | - | - |
|  |  | 40-59 | - | - |
|  |  | 60-74 | - | -2.94 (-7.48; 1.82) |
|  |  | Total | - | -1.72 (-6.04; 2.80) |
| Ecuador | 1997-2013 | 25-39 | - | - |
|  |  | 40-59 | -0.13 (-4.84; 4.81) | 2.11 (-2.23; 6.65) |
|  |  | 60-74 | 0.75 (-2.92; 4.57) | 2.01 (-0.88; 4.98) |
|  |  | Total | 0.49 (-2.38; 3.45) | 1.83 (-0.33; 4.03) |
| El Salvador | 1997-2013 | 25-39 | - | - |
|  |  | 40-59 | - | - |
|  |  | 60-74 | 1.47 (-3.79; 7.03) | -0.76 (-4.86; 3.51) |
|  |  | Total | -1.20 (-5.59; 3.39) | 1.89 (-2.44; 6.41) |
| Guatemala | 2000-2013 | 25-39 | - | - |
|  |  | 40-59 | - | - |
|  |  | 60-74 | 6.75 (-1.07; 15.18) | 8.37 (2.02; 15.13) |
|  |  | Total | 5.74 (-0.69; 12.58) | 8.77 (3.06; 14.81) |
| Mexico | 1998-2013 | 25-39 | 3.60 (-2.68; 10.29) | 2.73 (-0.91; 6.51) |
|  |  | 40-59 | 0.75 (-1.58; 3.14) | 0.63 (-0.48; 1.74) |
|  |  | 60-74 | 0.68 (-0.86; 2.24) | 0.68 (-0.05; 1.41) |
|  |  | Total | 0.71 (-0.30; 1.74) | 0.73 (0.20; 1.27) |
| Nicaragua | 1997-2013 | 25-39 | - | - |
|  |  | 40-59 | - | - |
|  |  | 60-74 | - | - |
|  |  | Total | - | 4.07 (0.07; 8.24) |
| Panama | 1998-2013 | 25-39 | - | - |
|  |  | 40-59 | - | - |
|  |  | 60-74 | - | - |
|  |  | Total | - | -0.20 (-4.40; 4.18) |
| Paraguay | 1996-2013 | 25-39 | - | - |
|  |  | 40-59 | - | - |
|  |  | 60-74 | - | - |
|  |  | Total | - | 1.82 (-1.95; 5.73) |
| Peru | 1999-2013 | 25-39 | - | - |
|  |  | 40-59 | 3.87 (-3.58; 11.89) | -2.17 (-4.52; 0.23) |
|  |  | 60-74 | 0.49 (-3.45; 4.60) | 0.56 (-1.03; 2.18) |
|  |  | Total | 1.51 (-1.83; 4.97) | -0.33 (-1.39; 0.74) |
| Uruguay | 1997-2010 | 25-39 | - | - |
|  |  | 40-59 | - | - |
|  |  | 60-74 | -1.36 (-5.12; 2.55) | -0.22 (-6.42; 6.38) |
|  |  | Total | -0.82 (-7.24; 6.04) | -1.86 (-7.70; 4.35) |
| Venezuela | 1996-2013 | 25-39 | - | - |
|  |  | 40-59 | -1.06 (-3.55; 1.49) | 0.06 (-1.51; 1.66) |
|  |  | 60-74 | 0.52 (-1.79; 2.88) | 1.12 (-0.47; 2.74) |
|  |  | Total | -0.07 (-1.77; 1.66) | 0.66 (-0.35; 1.69) |

## DISCUSSION

The incidence of thyroid cancer occurred more frequently in the age group of 40 to 59 years in Costa Rica and Quito (Ecuador), while in Cali (Colombia), Goiania (Brazil) and Valdivia (Chile) there was a higher incidence in the age group over 60 years. The highest rates were observed in women, found similar to other studies (8, 17).

Increased incidence trends were detected for the women and the stability trends were observed in Cali, Goiania, Quito and Valdivia for men in the age group above 60 years. The increase in incidence trends, for both sexes, is also observed in developed and developing countries, such as Denmark (male: AAPC = 3.2%; p< 0.05 and female: AAPC = 3.6%; p < 0.05); United States (male: AAPC = 3.1%; CI 95%: 2.7−3.5 and female: AAPC = 3.7%; CI 95%: 3.3-4.1) and China with AAPC =22.86% (CI 95%: 19.2-26.7), similar to both sexes (18–20).

The main risk factor associated with thyroid cancer is ionizing radiation because this gland is radiosensitive in young age and is in a position that allows for greater uptake of radiation. This increased incidence may be related to increased individual radiation dose observed in recent decades, through medical and dental diagnostic procedures, which has greater impact when exposure occurs during childhood (21, 22).

The factors that contribute to the temporal trends and geographic variations include the prevalence of obesity and diabetes among the countries (23, 24). It can be verified that in Latin America and the Caribbean, 16% of men and 20% of women were obese (25). Another factor is the enrichment of iodine in the diet, whose risk for thyroid cancer may differ based on the availability of iodine, is excess or scarcity (3, 26). By analyzing the incidence of thyroid cancer in São Paulo (Brazil) and the United States it was found that the differences in the nutritional status of iodine among populations may have affected the observed incidence patterns (27).

Other possible risk factors associated with women are sex hormones in interaction with the Thyroid-Stimulating Hormone (TSH), which can play a critical role in the development of thyroid malignancy, as well as advanced age in menopause and the greater parity (28, 29). In men, it was found a positive association with the story of goiter, thyroid nodules and family history of cancer (30). However, the consumption of tobacco and alcohol were not associated with increased risk (31, 32).

However, authors discuss whether there is a real increase of thyroid cancer or dealing with an epidemic of diagnoses, due to the heterogeneous pattern between a rising incidence and mortality rates (10, 33). This increased incidence is assigned to detection of subclinical disease and non-lethal, for better access to the health system, incidental detection of the image and more frequent biopsy (34, 35). Sierra et al. (5) showed that the increased incidence of thyroid cancer in Argentina, Brazil, Chile and Costa Rica was driven primarily by the increased incidence of papillary subtype.

The distinctive thyroid cancer, follicular and papillary subtypes, which represents the highest percentage of all thyroid cancers and is responsible for the increasing incidence of the disease, presents very good prognosis and low mortality (36). The estimated survival rate of 5 years for thyroid cancer is excellent when identified. The age has a strong effect on the disease-specific survival because it decreases with the increasing of age. Furthermore, males were associated with a significantly worse survival rate among patients with regional disease (37).

In most of the countries of Latin America there was stability in the mortality trend. Corroborating with the finds in China, there was stability in mortality from thyroid cancer (AAPC = 2.05%; CI 95%:−1.7-6.0), for both sexes, and Central Serbia, male AAPC= 2.4% (CI 95%: −0.5-5.5) and female AAPC=−1.3% (CI 95%: −4.4-1.9) (19, 38). In the United States, in the 1994-2013 period, the mortality increased 0.9% per year (CI 95%: 0.7 −1.5), for both sexes (20).

Only Argentina, Brazil, Chile, Cuba and Nicaragua in Latin America are implementing policies, strategies or action plans specifically for cancer. Countries such as Belize, Ecuador, Mexico, Nicaragua and Panama are advancing in this regard, while several countries in the region have an absence (or lack) of relevant information that would make possible their monitoring strategies for cancer control (39).

In regard to the quality of the data, the absence of population-based registers with historical series made it impossible for the inclusion of other data. To mortality, differences were found in the comprehensiveness and completeness of the 17 countries studied, ranging from 55% in the Dominican Republic completeness and 90% in Argentina, Chile, Costa Rica, Mexico, Uruguayand Venezuela. In addition, the percentage of ill-defined deaths ranged from 5% (Costa Rica and Mexico) to 24% (El Salvador) (40). Despite these limitations, the data have been validated by international organizations and can be used to describe the mortality in Latin American countries (41–43).

This study analyzed the incidence of thyroid cancer in four cities and one country of Latin America (Costa Rica) and the mortality trends for thyroid cancer in 16 countries of Latin America. Both incidences as mortality showed differences, with an increase in women of Cali (Colombia), Costa Rica, Goiania (Brazil) and Quito (Ecuador). There was stability in mortality trends in most countries of Latin America.

## AUTHOR CONTRIBUTIONS

Conceptualization: DS MC IR.

Data curation: FF IR.

Formal analysis: FF IR.

Funding acquisition: FF IR.

Investigation: FF DS MC IR.

Methodology: DS MC IR.

Project administration: IR.

Resources: FF DS MC IR.

Software: DS MC IR.

Supervision: DS IR.

Validation: DS MC IR.

Visualization: FF DS MC IR.

Writing ± original draft: FF.

Writing ± review & editing: FF DS MC IR.

